# Preferential apical infection of intestinal cell monolayers by SARS-CoV-2 is associated with damage to cellular barrier integrity: Implications for the physiopathology of COVID-19

**DOI:** 10.1101/2024.01.08.574642

**Authors:** Clémence Garrec, Jeffrey Arrindell, Jonatane Andrieu, Benoit Desnues, Jean-Louis Mege, Ikram Omar Osman, Christian A. Devaux

## Abstract

SARS-CoV-2 can infect different organs, including the intestine. In Caco-2 intestinal cell line, SARS-CoV-2 modulates the ACE2 receptor expression and affects the expression of molecules involved in intercellular junctions. To further explore the possibility that the intestinal epithelium serves as an alternative infection route for SARS-CoV-2, we used a model of polarised intestinal cell monolayers grown on the polycarbonate membrane of Transwell inserts, inoculated with the virus either in the upper or lower chamber of culture. In both polarised Caco-2 cell monolayers and co-culture Caco-2/HT29 cell monolayer, apical SARS-CoV-2 inoculation was found to be much more effective in establishing infection than basolateral inoculation. In addition, apical SARS-CoV-2 infection triggers monolayer degeneration, as shown by histological examination, measurement of trans-epithelial electronic resistance, and cell adhesion molecule expression. During this process, the infectious viruses reach the lower chamber, suggesting either a transcytosis mechanism from the apical side to the basolateral side of cells, a paracellular trafficking of the virus after damage to intercellular junctions in the epithelial barrier, or both. Taken together, these data highlight a preferential tropism of SARS-CoV-2 for the apical side of the human intestinal tract and suggests that infection via the intestinal lumen leads to a systemic infection.

## Introduction

SARS-CoV-2 was identified in 2019 as a novel emergent Sarbecovirus causing severe human respiratory infection called coronavirus disease 2019 (COVID-19). As of 12 April 2023, SARS-CoV-2 had been responsible for more than 760 million cumulative cases (762,791,152 cases) and more than six million deaths (6,897,025 deaths) (**WHO Coronavirus Dashboard;** accessed 19 April 2023). Direct or indirect contact with respiratory droplets of SARS-CoV-2 from a source patient is considered to be the main route for human-to-human virus transmission. Common symptoms associated with SARS-CoV-2 infection are fever, cough, and shortness of breath. In addition, gastrointestinal symptoms such as abdominal pain, nausea, vomiting, and diarrhoea, are among the most commonly reported extrapulmonary clinical features of COVID-19 and have been noted in 20% to 50% of patients, sometimes preceding the development of respiratory disease[1–3]. A multicentre study in China found that patients with diarrhoea more frequently showed severe symptoms of pneumonia than patients without diarrhoea (53% versus 19%)[4]. Diarrhoea is thought to be induced either by dysbiosis in the gut microbiota, the release of toxins, enterocyte damage leading to malabsorption and/or inflammation followed by intestinal fluid osmotic dysfunction, and/or electrolyte secretion by activation of the enteric nervous system. SARS-CoV-2 has been observed to productively infect human enterocytes in human small intestinal organoids (hSIOs or “mini-gut”), with significant titres of infectious viral particles being detected, although shedding of the Omicron strain was found to be severely reduced compared with the Wuhan and Delta strains[5,6]. SARS-CoV-2 RNA, as well as live virus have been found in the stools of COVID-19 patients[7,8] and both viral nucleocapsid proteins and viral particles have been found in infected intestinal biopsies from COVID-19 patients[9]. From a growing pool of evidence supporting the hypothesis that persistent SARS-CoV-2 could sustain inflammation and blood clots in patients categorised as having “long COVID” (patients for whom symptoms linger beyond 12 weeks/84-days after an acute infection), the idea recently emerged that the gut epithelium might be a long-term reservoir of lingering virus, driving long COVID[10–15]. It was recently published that in the case of long COVID without any detectable viral particles in the respiratory tract, 12.7% of patients had SARS-CoV-2 in their stools at 120 days post-infection and 3.8% still excreted SARS-CoV-2 at 210 days post-infection[16]. In addition, viable virus was found in the appendices of two patients with long COVID symptoms at 175 days post-infection and 462 days post-infection, respectively. This was the first study to detect a viable virus for so long in the digestive tract[17]. These results drastically change our understanding of the role of enterocyte infection by SARS-CoV-2 in the pathophysiology of COVID-19[3,18].

Using animal models to investigate the pathophysiology of gastrointestinal SARS-CoV-2 infection, it has been reported that the intragastric inoculation of nonhuman primates with SARS-CoV-2 resulted in the productive infection of digestive tissues and inflammation in both lung and intestinal tissues[19]. This route of inoculation induces the production of inflammatory cytokines and immunohistochemistry showed decreased Ki67, increased cleaved caspase 3, and decreased numbers of mucin-containing goblet cells, suggesting that the inflammation process impaired the gastrointestinal barrier. The intestinal epithelium is composed of a monolayer of self-renewing intestinal epithelial cells linked together by intercellular junctions resulting from interaction between cell adhesion molecules (CAMs) necessary for the integrity of the intestinal barrier[20]. The epithelial cells layer is polarised, organised into an apical domain which faces the lumen of the intestine and a basolateral domain divided into a basal domain facing the basal membrane and a lateral domain facing the neighbouring cells. This structural tissue integrity is preserved through interactions involving tight junctions (TJs), adherents junctions (AJs), desmosomes, and gap junctions[21,22]. Epithelial cell polarity allows a balanced communication between the intestinal lumen and the tissues, and provides a defence function against gastrointestinal pathogens[23]. We recently reported a significant decrease in the expression of CDH1/E-cadherin by Caco-2 cells when they were infected with SARS-CoV-2, as well as an increase in the production of the soluble form of E-cadherin (sE-cad)[24]. As well, we found that infection of Caco-2 cells by SARS-CoV-2 induces the dysregulation of other adhesion molecules (Occludin, JAMA-A, Zonulin, Connexin-43 and PECAM-1). In agreement with our study and using a model of human-induced pluripotent stem cell-derived small intestinal epithelial cells (iPSC-SIECs), it was observed, that SARS-CoV-2 infection decreased expression levels of tight junction markers, ZO-3 and CLDN1, and the transepithelial electrical resistance (TEER), which evaluates the integrity of tight junction dynamics[25]. In addition, in the animal model of K18-hACE2 transgenic mice expressing human ACE2 infected with SARS-CoV-2 via intranasal administration, we found extensive necrosis and inflammation of the lamina propria of intestinal villi associated with cell damage and increased intestinal permeability[24].

One important question that remains to be answered concerns the route of infection of intestinal cells. This could be either a direct infection at the apical pole after the virus passes through the oral cavity, the oesophagus, and the digestive tract, or an infection at the basal pole after a pulmonary infection and passage in the blood circulation, allowing the virus to have access to the basal regions of the intestinal tissue. Although most viruses would probably die in the harsh acid environment of the stomach, it remains possible that the saliva and secretions could carry the virus into the digestive tract where viral replication may be sustained in epithelial cells[1], as already demonstrated for many enteroviruses. Moreover, it was reported that although the virus is inactivated by gastric pH of <2.5, it is much less affected by pH between 3.5 and 6.0 that fits gastric pH variations depending whether the individual is in fasting or feeding state, respectively[26]. Finally, a recent study provided evidence for the transmission of SARS-CoV-2 via people’s hands through frequently-touched surfaces at home[27]. The present study, aimed at distinguishing the different possible routes of infection of intestinal cells and their sensitivities to polarised infection (apical and/or basal) by SARS-CoV-2, revealed a preferential tropism of SARS-CoV-2 for the apical surface of the human intestinal cells, as well as damage to the integrity of the epithelial cell monolayer.

## Materials and methods

### Cells culture

Caco-2 (ATCC® HTB-37 ™), isolated from a human colorectal adenocarcinoma able to exhibit spontaneous epithelial differentiation in vitro, and HT29 (ATCC® HTB-38 ™), another human tumour epithelial cell line of intestinal origin producing mucin, were cultured in Dulbecco’s modified Eagle medium with high glucose content (DMEM-F12, Gibco, Waltham, MA, USA) supplemented with 10% foetal bovine serum (FBS, Gibco). Vero-E6 (ATCC® CRL-1586 ™), a renal epithelial cell line of Simian origin isolated from Chlorocebus sabaeus (African green monkey), was cultured in Minimum Essential Medium (MEM, Gibco; Invitrogen) containing 4% FCS and 1% L-Glutamine (L-Gln; Invitrogen). These adherent cells were cultured to confluence for five days at 37 °C in an atmosphere containing 5% CO_2_.

### Cell culture in Transwell inserts

Caco-2 cell cultures (at a cell density of 10^5^ cells/cm2) or Caco-2/HT29 co-cultures (at a cell density of 10^5^ cells.cm^-2^/10^4^ cells.cm^-2^) were grown for 21 days in 12-well Transwell inserts (pore size 0.4 μm) (Corning) in the presence of DMEM-F12 medium supplemented with 10% FBS. A change of medium was performed every 2 days. The cells were considered to be confluent and polarised on the polycarbonate membrane of Transwell inserts when a stable measurement of the transepithelial electrical resistance (TEER) using the Millicell ERS-2 Volt-Ohm meter (EMD Millipore) reached 500∼680 Ω.cm^2^, as previously described[28].

### Virus production

SARS-CoV-2 (Wuhan strain: IHUMI-3) kindly provided by Prof. Bernard La Scola, was previously isolated from fluid collected in a nasopharyngeal swab specimen[29]. Vero-E6 cells were used to culture the viral isolate. After three passages and an almost complete cytopathic effect, the supernatant of each viral culture was collected, centrifuged at 3000 × g for 10 minutes at 4 °C, and then filtered through a 0.22 μm membrane. The filtrate was supplemented with 10% FBS and 1% 2-[4-(2-96 hydroxyethyl)piperazin-1-yl]ethanesulfonic acid (HEPES) and stored at -80 °C, to keep the viral stock of SARS-CoV-2. It is worth noting that upon passing in Vero-E6 cells, SARS-CoV-2 is apparently under strong selection pressure to acquire adaptive mutations in the multibasic S1/S2 cleavage site. This was evidenced by plaque assay showing remarkable plaque heterogeneity with mutants harboring 10 aa deletion or an Arg682Gln substitution[30]. Although our viral stock has not been subjected to resequencing before use, IHU-MI3 strain is closely related to SARS-CoV-2 Wuhan IHUMI-3 reference strain[31–33].

### Infection of cell monolayers in Transwell inserts

After 21 days of culture and a stable TEER of 500∼700 Ω.cm2, polarised Caco-2 cells or polarised Caco-2/HT29 co-cultures were infected with an inoculum of SARS-CoV-2 IHUMI-3 at an MOI (Multiplicity of Infection) of 0.05 suspended in DMEM-F12 culture medium (10% FBS), either on the apical (upper chamber) or basolateral side (lower chamber). At 4h, 8h, 16h, 24h and 48h post infection, the transepithelial electrical resistance (TEER) was measured and the supernatants from the upper and lower chambers of the Transwell inserts were recovered in aliquots then frozen at -80 °C until use.

### RNA extraction and qRT-PCR

SARS-CoV-2 RNA was extracted from 100µL of cell culture supernatants from the upper and lower chambers of the Transwell inserts recovered at 4h, 8h, 16h, 24h and 48h post-infection, using the QIAamp 96 Virus kit QIAcube HT (Qiagen). To detect SARS-CoV-2 RNA, real-time RT-PCRs were performed using a set of SARS-CoV-2 N-gene primers (Fwd: 5’-GACCCCAAAATCAGCGAAAT-3’, Rev: 5’-TCTGGTTACTGCCAGTTGAATTCG-3’; 5’ probes FAM-ACCCCGCATTACGTTTGGTGGACC-QSY 3’) and Quantitative Superscript III Platinum One-step RT-qPCR systems with the ROX kit (Invitrogen), according to the manufacturer’s recommendations. The amplification cycles were carried out on a Light Cycler 480 (Roche Diagnostics). The results of viral qRT-PCR were expressed as Ct or as Relative quantification (2^-ýCT^), where ΔCt = Ct (T0) – Ct (T_post-infection_).

To quantify the expression of several genes in the cellular models studied, RNA was extracted from the Caco-2 cell lines using a RNeasy Mini Kit (QIAGEN SA) with a DNase I step to eliminate DNA contaminants. The quantity and quality of the RNAs was evaluated using a Nanodrop 1000 spectrophotometer (Thermo Science). The first strand cDNA was obtained using oligo(dT) primers and Moloney murine leukaemia virus-reverse transcriptase (Life Technologies), using 100 ng of purified RNA. The qPCR experiments were performed using specific oligonucleotide primers previously described[24] and hot-start polymerase (SYBR Green Fast Master Mix; Roche Diagnostics). The amplification cycles were performed using a C1000 Touch Thermal cycler (Biorad). The results of qRT-PCR were normalised using the housekeeping gene β-actin, and expressed as relative expression (22^-ýCT^), where ΔCt = Ct (Target gene) – Ct (Actin).

### TCID_50_ quantification of viral production

Viral release was also evaluated by tissue culture infectious dose 50 (TCID_50_). For this purpose, 96-well plates containing 1×10^5^ Vero E6 cell/well were prepared one day in advance and kept overnight in a 5% CO2 atmosphere at 37 °C. Culture supernatants from Caco-2 cells collected at 4h, 8h, 16h, 24h, 48h, and 72h were thawed and serially diluted at base ten and inoculated in Vero E6 cells (eight replicates per dilution to a 10-10 dilution). The plates were kept at 37 °C in a 5% CO2 atmosphere for seven days. Seven days post infection the presence of cytopathic effects was evaluated. The TCID_50_ was calculated according to the Spearman and Kärber algorithm[34].

### Confocal microscopy analysis

Monolayer polycarbonate membranes with polarised cells infected or not with SARS-CoV-2 (MOI: 0,05) were fixed with 3% paraformaldehyde (PFA) for 15 minutes at 4 °C then cut with a scalpel and preserved in formaldehyde. Part of the polycarbonate membranes with cells was dehydrated and clarified with xylene by an automaton (TP1020, Leica Biosystems, Heidelberg, Germany). Cell preparations were manually embedded in paraffin so that sagittal sections could be obtained. The paraffin blocks containing the polycarbonate membranes surmounted by the cell monolayer were cut with a Histocore Rotary microtome (Leica Biosystems). The sections were then deparaffinised, stained with Hemalun, Phloxine and Safran (HPS), as well with Periodic Acid-Schiff (PAS) for the mucin detection in the Caco-2/HT29 coculture and observed with a Zeiss LSM800 confocal microscope. The other part of the polycarbonate membranes surmounted by the cell monolayer was included to obtain sagittal sections, in OCT (Optimal Cutting Temperature) embedding medium. Cryosections were then performed using a Cryostat CM1860 (Leica Biosystems) at a temperature below -20 °C and placed on slides. The cryostat sections were fixed for 10 minutes with 96% acetone then ventilated for 20 minutes, followed by incubation with an anti-E-cadherin mouse mAb (4A2C7, Life Technologies, France) directed against the cytoplasmic domain of E-cad and a monoclonal anti-Occludin mouse mAb (331500, Life Technologies SA) and revealed using an anti-mouse IgG (H+L) secondary antibody (Alexa Fluor 555) (Life Technologies). For the SARS-CoV-2 staining a polyclonal rabbit anti-SARS Coronavirus Spike protein (PA1-41142, Thermo Fisher, France) and revealed using an anti-rabbit IgG (H+L) secondary antibody (Alexa Fluor 647) (Life Technologies,France). 4’,6’-diamino-2-fenil-indol (DAPI) (1:25000, Life Technologies) and Phalloidin (Alexa 488) (1:500, OZYME) were used respectively for nucleus and filamentous actin staining. Fluorescence and image acquisition were performed using a confocal Zeiss LSM 800 microscope (Carl ZeissMicroscopy, Germany).

### Electron microscopy analysis

The monolayer of Caco-2 cells polarised in a Transwell 12 culture insert for 21 days and infected with SARS-CoV-2 at an MOI of 0.05 was recovered and then fixed with 2.5% glutaraldehyde. Sagittal sections of the polarised cell monolayer were embedded in resin using the PELCO BioWavePro+ (Ted Pella Inc., USA). Sections were washed with 0.2M sucrose/0.1M sodium cacodylate and post-fixed with 1% OsO4 diluted in 0.2M potassium hexacyanoferrate (III)/sodium cacodylate buffer 0.1M. After washing with distilled water, the samples were gradually dehydrated by successive baths containing 30% to 100% ethanol. Substitution with Epon resin was achieved by incubations with 25% to 100% Epon resin, and the samples were placed in a polymerisation chamber. The microwave curing of the resin was carried out for a total of two hours. After hardening, the resin blocks were extracted from the Eppendorfs and manually trimmed with a razor blade. The resin blocks were placed in an EM UC7 ultramicrotome (Leica Biosystems), cut into pyramids, and ultrathin 300 nm sections were cut and placed on HR25 300 Mesh Copper/Rhodium (TAAB) grids. The sections were contrasted with uranyl acetate and lead citrate. Grids were attached with double-side tape to a glass slide and platinum-coated at 10 mA for 20 seconds with a MC1000 sputter coater (Hitachi High-Technologies, Japan). Electron micrographs were obtained on a SU5000 SEM (Hitachi High-Technologies, Japan) operated in high-vacuum at 7 kV accelerating voltage and observation mode (spot size 30) with BSE detector. As for the transmission electron analysis, after contrast with uranyl acetate and lead citrate, was carried out using a Tecnai G20 Cryo transmission electron microscope (FEI, 200 keV).

### Statistical analysis

Analysis of variance (ANOVA) was performed using the GraphPad-Prism software (version 9.0). Data were analysed using a one or a two-way ANOVA with the Holm-Sidak multiple post-hoc test for group comparison. A *P*-value <0.05 was considered statistically significant. The results are presented as the mean with standard error of the mean (SEM).

## Results

### Characterisation of the cellular model

A cellular model regularly used to mimic enterocytes is the human colon adenocarcinoma Caco-2 cell line, which is susceptible to SARS-CoV-2 infection and intense viral replication without cytopathic effect. Caco-2 cells cultured in Transwell inserts can undergo spontaneous differentiation, display morphological and functional features of enterocytes, and form an intact permeability barrier, which could be reflected by heightened transepithelial electronic resistance (TEER), and the formation of polarised cell adhesion architecture.

To carry out our study, we first had to select a cellular model of polarisable epithelial cells sensitive to infection by SARS-CoV-2. As we had previously demonstrated the effect of SARS-CoV-2 infection on a wide range of genes in CaCo-2 cells, we chose this cell line as a model. For an overview of this preliminary screening, we have summarized in Figure 1A the gene expression in virus-free Caco-2 cells whose basal expression was affected by the presence of the virus.

**Figure 1.**
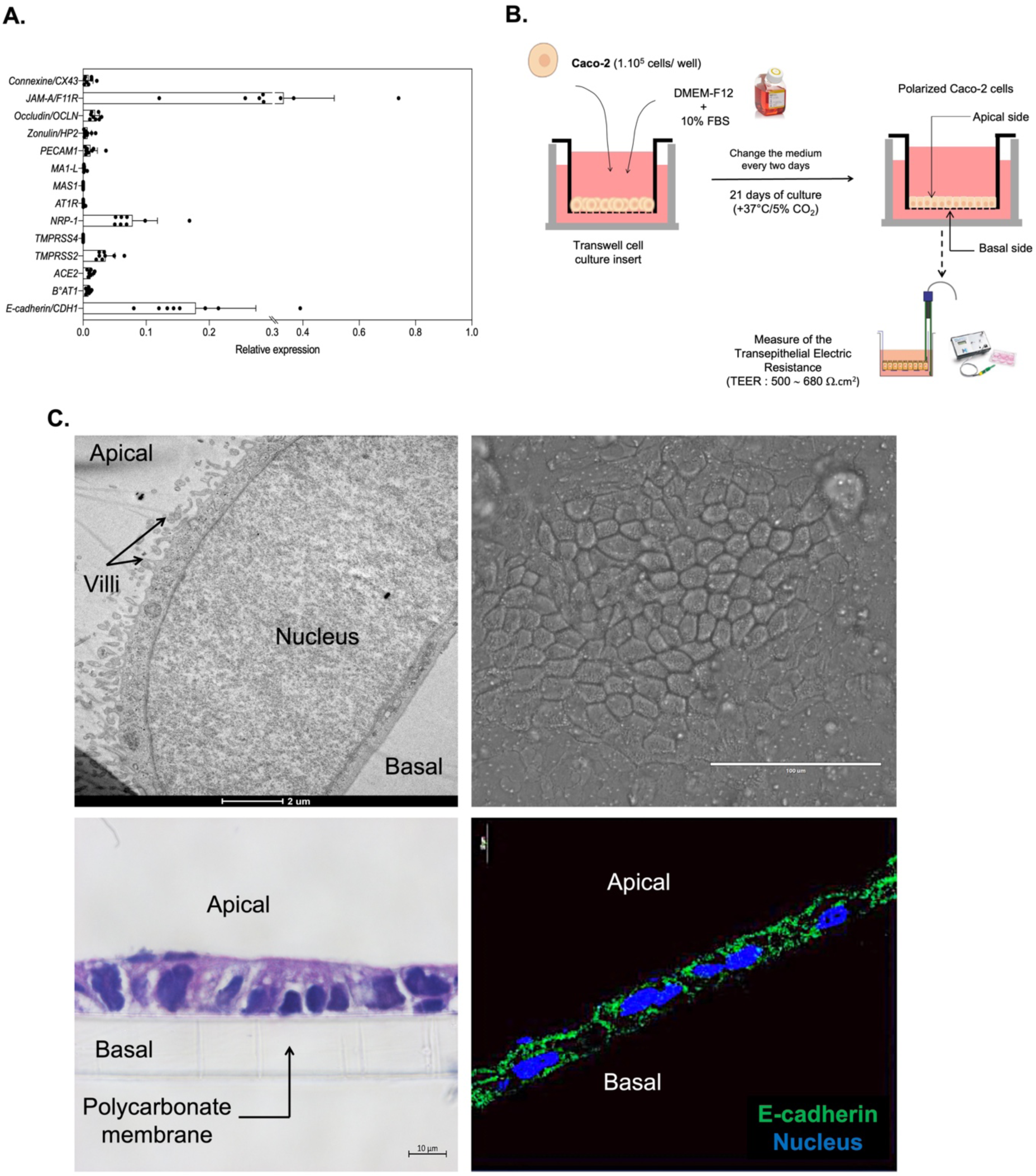
Characterization of the cellular model to study the effect of SARS-CoV-2 inoculation on the structure and the permeability of epithelial monolayer. **(A)** Basal expression of the the mRNA coding proteins involved in intercellular junctions and SARS-Cov-2 infection in in virus-free unpolarized Caco-2 cells. **(B)** Schematic flow of the analysis: After 21 days of culture in Transwell insert, the Caco-2 cells adopted an epithelial monolayer resembling an epithelial permeability barrier. The electronic trans-epithelial resistance (TEER) was measured during culture and was considered stable when it reached ∼500-700 Ω.cm^2^. **(C)** Heatmap of mRNA expression of a panel of genes in virus-free Caco-2cells. The results (n=8) are expressed as RE where RE=2^(−ΔCT)^. the Heatmap was generated using the GraphPad (9.0) software. (C) Microscopy analysis of the Caco-2 cells monolayer. Transmission electron microscopy (upper/left panel; scale bare: 2μm); Bright fields (upper/right panel; scale bare: 100 μm) observation; HPS coloration (Lower/left panel; scale bare: 10 μm); Cellular localization/distribution of E-cad protein (red), actin (green) and nucleus (blue) (Lower/right panel; scale bare : 10 μm).

After 21 days of culture in Transwell inserts (**Fig. 1B**), Caco-2 cells adopted an architecture of adherent polarised cells and formed an epithelial monolayer resembling an epithelial permeability barrier (**Fig. 1C**). The integrity of the permeability barrier was evaluated by measuring the trans-epithelial electronic resistance (TEER) and a stable TEER of ∼500–700 Ω.cm2 was considered a strong indicator of the integrity of the epithelial permeability barrier.

### Expression of cell adhesion proteins after SARS-CoV-2 inoculation at the apical or basolateral membrane of Caco-2 cells

To analyse the effect of SARS-CoV-2 inoculation on the structure and permeability of the epithelial monolayer, different type of experiments, summarised in (Figure 2A), were performed. The virus-free Caco-2 cell monolayers were then apically or basolaterally inoculated with SARS-CoV-2 at an MOI of 0.05. The viral input was not washed out to avoid mechanical damage to the cell monolayer, and the TEER was measured at different post-infection times.

**Figure 2.**
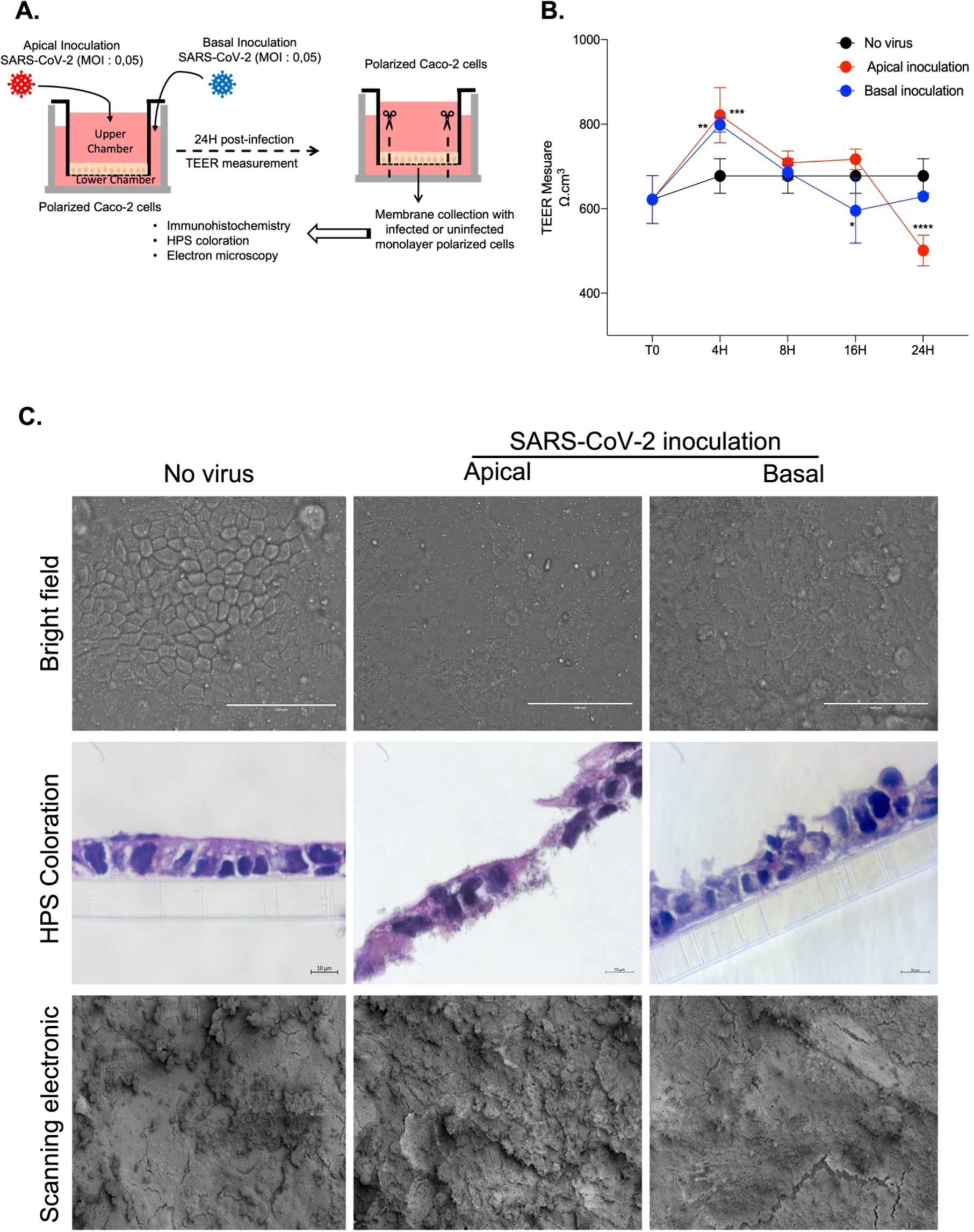
Impact of SARS-CoV-2 inoculation on the structure and permeability of the epithelial monolayer of Caco-2 cells. (**A**) Schematic flow of infection experiment: Caco-2 cells monolayer was inoculated with the virus at apical (upper chamber) or basal side (lower chamber). At different time of infection, the TEER was measured. At 48h post-infection, the polycarbonate membrane to which the monolayer cells adhere was collected for further microscopy analysis. (**B**) TEER measure (n=3) at 4h, 8h, 16h, 24h, 48h post-inoculation to evaluate the permeability of the Caco-2 cells monolayer. (**C**) Microscopy analysis of the Caco-2 cells inoculated with virus at apical side or basolateral side. Bright fields (upper panel, scale bare: 100 μm) observation; HPS coloration (Middle panel, scale bare : 10 μm); Scanning electron microscopy (Lower panel; scale bare: 100μm).

As result and compared to the virus-free polarized cells, at eight hours post-inoculation, ohmic resistance start to decrease (**Fig. 2B**) and become statistically significant at 16h and 24h, with a mean TEER value of 443 Ω.cm^2^ at t=24 h after apical inoculation and of 556 Ω.cm^2^ at t=24 h after basolateral inoculation. This decrease in TEER, particularly when the virus is inoculated on the apical side, leads to an architectural modification of the Caco-2 cell monolayer, as shown by light microscopy (bright field) and scanning electron microscopy (**Fig. 2C** upper panel and lower panel, respectively) of the cells 48 hours after inoculation. Additionally, the confocal microscopy analysis (HPS colouration) (**Fig. 2C** middle panel) indicated that the monolayers were badly damaged, while the cells tended to detach from the polycarbonate support.

To confirm this structural damage, the expression of intercellular adhesion proteins in virus-free Caco-2 cell monolayers and cells exposed to infectious SARS-CoV-2 by inoculation either at the apical or basolateral side were observed by confocal immunofluorescence analysis. As shown in **Fig. 3A**, the virus-free Caco-2 cell monolayers show high expression of E-cadherin (E-cad), but when the monolayers were inoculated with SARS-CoV-2 via the apical side, at 24 hours post-infection the expression of E-cad on the monolayer was significantly reduced. This result suggests either an increase in soluble E-cadherin shedding or a reduced transcription of the CDH1 mRNA. In contrast, when the monolayers were inoculated with SARS-CoV-2 via the basolateral side, the intensity of E-cad protein expression was almost similar to that observed with virus-free Caco-2 cell monolayers, suggesting that the integrity of intercellular junction is maintained. The same results were obtained when cells were stained with an anti-occludin mAb (**Fig. 3B**). The fact that E-cad and occludin expression was reduced after SARS-CoV-2 inoculation via the apical side, correlated with the significant decrease of TEER and evidence of monolayer damage, while this phenomenon remained almost undetectable when SARS-CoV-2 was inoculated at the basolateral side of the monolayer.

**Figure 3.**
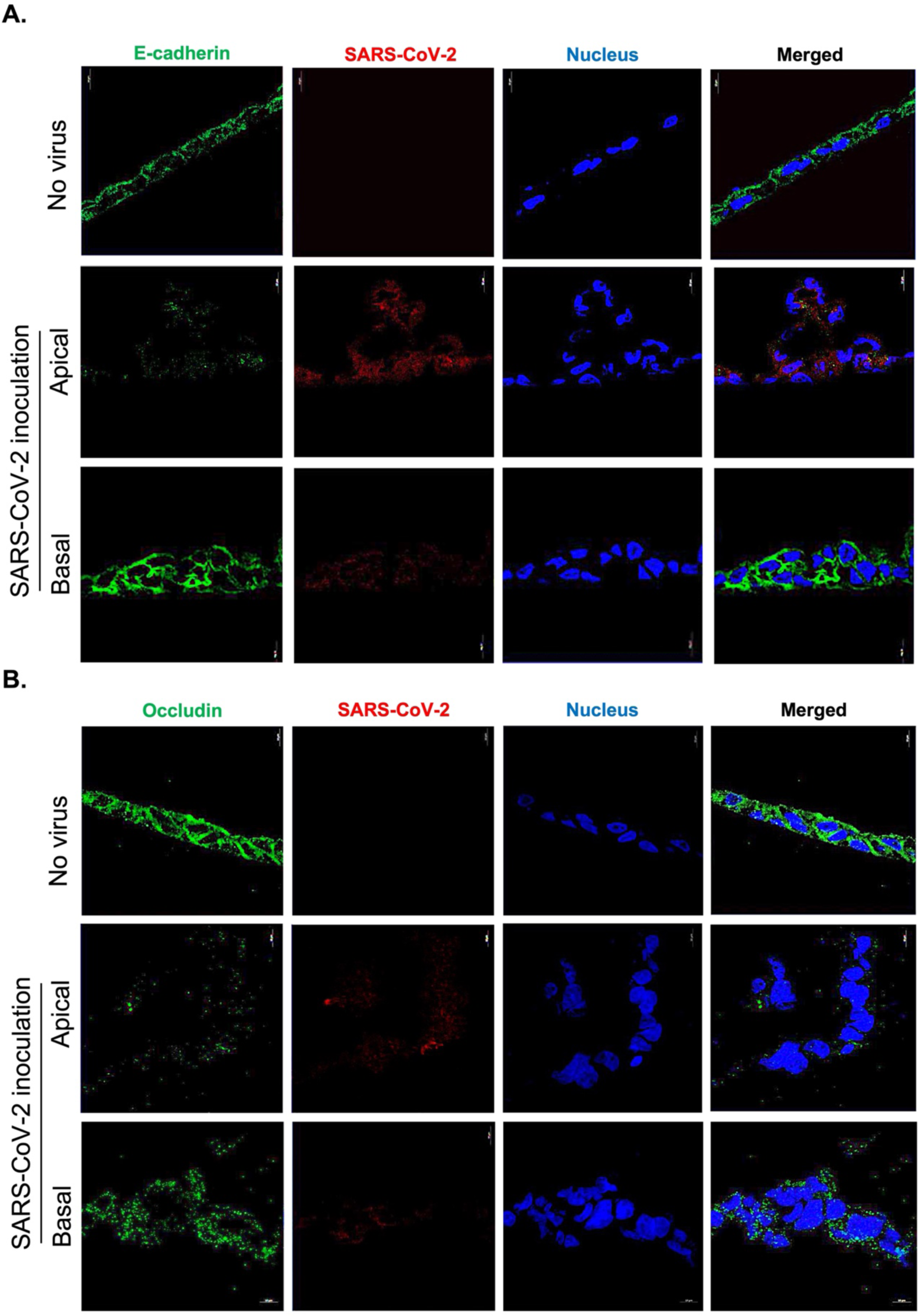
Modulation of cell surface expression of E-cadherin and Occludin in the Caco-2 cells monolayer inoculated or not with SARS-CoV-2 at 24 hours-post-inoculation. The panel presents monolayers cells free-virus versus SARS-CoV-2 apical and basolateral inoculated monolayer and evaluation of fluorescence corresponding to (**A**) the E-cad (green); (**B**) Occludin(green). The SARS-CoV-2 virus was stained in red and the nucleus of the cells in blue. Images were acquired using a confocal microscope (Zeiss LSM 800) with a 63X/1.4 oil objective (scale bar: 10 µm).

### SARS-CoV-2 inoculation and virus release in polarised intestinal epithelial cells

As a next step in the experiment, samples of cell-free media were harvested in the upper and lower chambers of the Transwell insert and the presence of cell-free virions (either viral input or progeny viruses) was evaluated in both chambers after SARS-CoV-2 inoculation either at the apical or basolateral side of the Caco-2 cell monolayers (**Fig. 4A**). When SARS-CoV-2 was inoculated at the apical side, a positive CT values were found (CT < 35) in samples collected from the upper chamber, which indicated the presence of the virus, while in the lower chamber the CT values remained negative (CT > 35) during the first 16 hours (**Fig. 4B**) post-inoculation. At 24 hours the CT values in the lower chamber decreased below 35 CT indicating the presence of the virion in the chamber. This occurrence can be explained by either a slow diffusion of virus particles from the upper chamber into the lower chamber as a consequence of a damaged monolayer barrier and paracellular trafficking or the production of progeny virions budding from the Caco-2 cells’ basolateral membrane after transcytosis (if the integrity of the epithelial barrier was sufficient to prevent virus diffusion).

**Figure 4.**
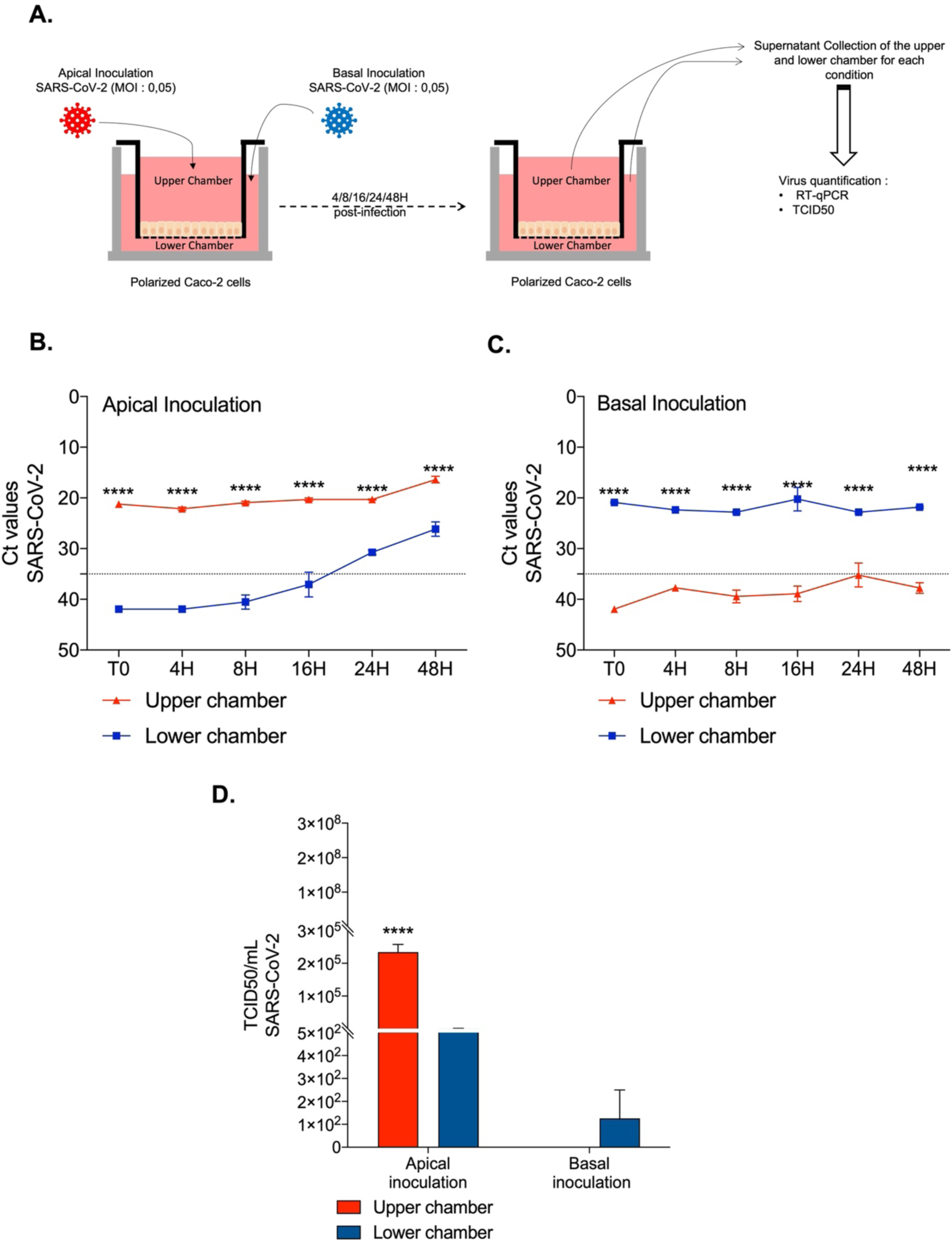
Kinetics of SARS-CoV-2 infection in Caco-2 monolayer cells. (**A**) Schematic flow of infection experiment: After viral inoculation at apical side (upper chamber) or basolateral side (lower chamber) in Caco-2 monolayer cells, the viral release was monitored over 48h hours (T=0,4h,8h,16h, 24h,48h) from the cell’s supernatant from upper and lower chamber. (**B)** and **C**) Quantification of viral release in the upper and lower chambers of the Caco-2 cell monolayer during apical and basolateral inoculation respectively by qRT-PCR (n=3). (**D**) Quantification of SARS-CoV-2 infectious particles released in the supernatant of Caco-2 cells monolayer after 48h post-infection. TDCI50 (n=3) was performed by inoculating supernatants from the upper or lower chambers from SARS-CoV-2 infected Caco-2 cell monolayers into Vero E6 cell culture. The cytopathic effect (CPE) was evaluated on Vero E6 cells at 7days after exposure to the supernatants. The TCID50/mL is calculated according to the Spearman and Kärber algorithm.

When SARS-CoV-2 was inoculated at the basolateral side of the monolayer, the low CT values (CT < 35) in the lower chamber indicated the presence of the virus (either viral input or progeny viruses). In contrast, the CT values in the upper chamber never reached values below 35, indicating that the virus was apparently contained in the lower chamber and that the integrity of the epithelial barrier was maintained (**Fig. 4C**).

To confirm this result, a more sensitive assay that consists of a titration of the TCID_50_ at 48 hours post-inoculation of each sample on Vero-E6 cells was performed. As shown in **Fig. 4D**, infective virus particles were bilaterally found into the media in the upper and lower chambers with apical side SARS-CoV-2 inoculation. In contrast, during the basolateral side virus inoculation, no infective virus particles were found in the upper chamber. Although the same concentrations of virus were used as input during both apical and basolateral inoculations, the TCID_50_ titre was 1 Log higher in the samples collected from the apical side after apical inoculation, suggesting the *de novo* synthesis of progeny virions and indicating that apical inoculation was much more effective in establishing a productive viral infection than basolateral inoculation.

In order to complete this analysis, we attempted to visualise viral particles either at the apical pole or at the basal pole, 72 hours after exposure of the monolayers of Caco-2 free-virus cells (**Fig. 5A**) to the virus. As can be seen in **Fig. 5B**, 72 hours after apical inoculation of virus on Caco-2 cell monolayers and compared to the basolateral inoculation (**Fig. 5C**), we can clearly distinguish SARS-CoV-2 particles at the apical membrane and villi (either residual viral input or progeny viruses). Despite the analysis of about a hundred histological sections by transmission electron microscopy, we did not observe any viral particles at the basolateral membrane (data not shown). Even if we cannot totally exclude the possibility that a process of transcytosis may exist, followed by virus budding at the basolateral membrane, these observations tend to indicate preferential budding at the apical pole and passage of the virus in the lower compartment only after loss of integrity of the Caco-2 cell monolayer.

**Figure 5.**
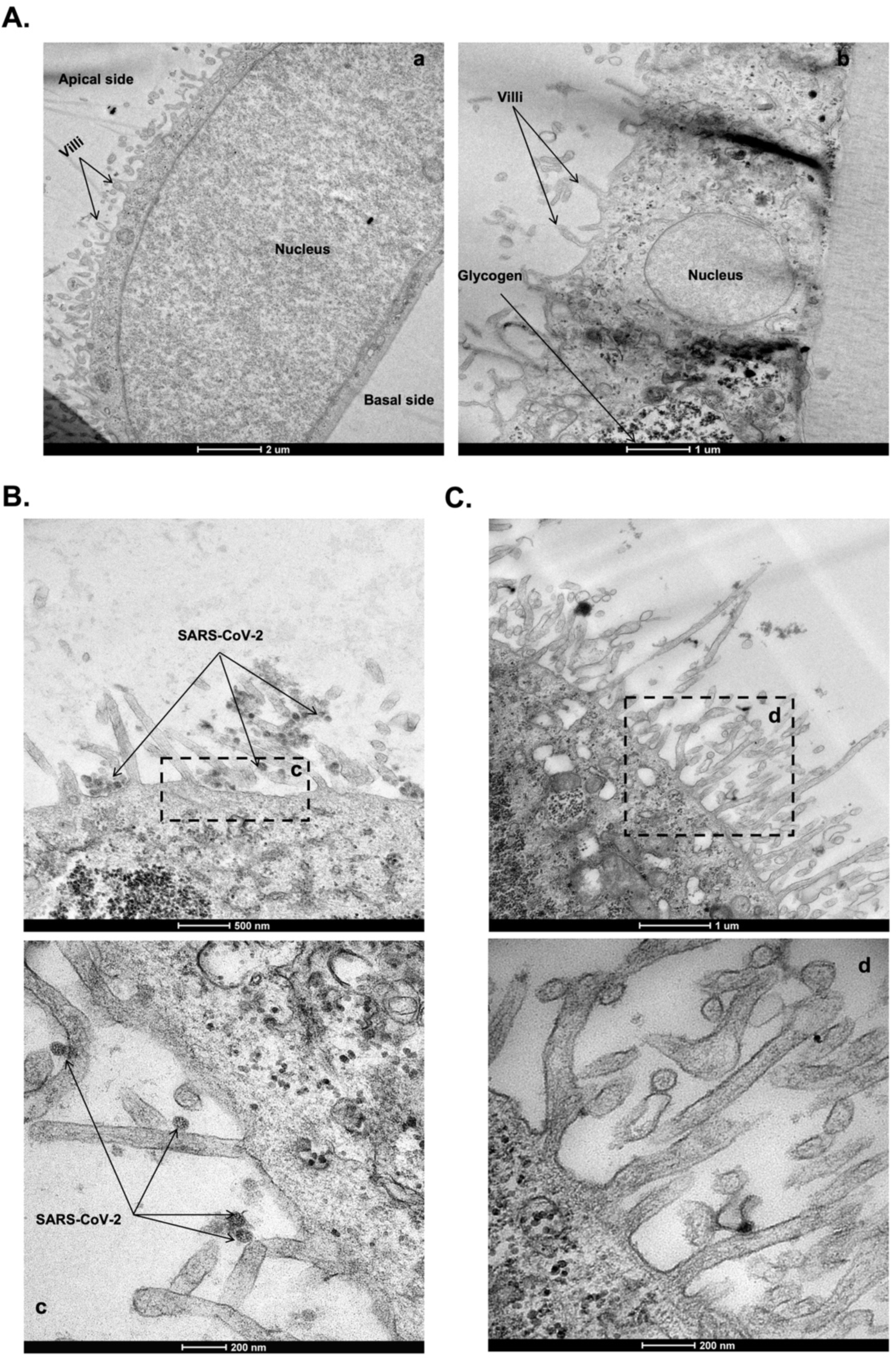
Transmission electron microscopy of SARS-CoV-2 infected Caco-2 monolayer cells. (**A**) Low (**a**) and Hight (**b**) magnification images of virus-free after 21 days of culture. (**B** and **C**) SARS-CoV-2 infected Caco-2 cells monolayer at 72 hours-post-inoculation. (**c**) Zoom in on the boxed region of (**B**) with SARS-CoV-2 particles located in the villi of the cell monolayer inoculated with the virus on the apical side. (**d**) Zoom-in boxed region Zoom in on the boxed region of (**C**) of the cell monolayer inoculated with the virus on the basolateral side.

These results align with the photonic and confocal microscopy showing that the Caco-2 cell monolayer appear disorganised and damaged when the virus is inoculated in the apical while it appears less damaged under basolateral inoculation, although TEER also decreased significantly on both sides of virus inoculation.

### SARS-CoV-2 inoculation and virus release in polarised intestinal epithelial cell monolayers composed of polarised Caco-2 cells and mucin-producing HT29 cells

Caco-2 cell culture monolayer systems are oversimplified and do not recapitulate the multiple cellular components, complex structure, and functions of the native intestine. In particular, they lack the mucin microenvironment considered as a typical characteristic of the intestinal barrier and playing a major role in the dynamic of viral infections including SARS-CoV-2. Thus, the lack of a mucus layer may facilitate SARS-CoV-2 binding to and entry into Caco-2 cells and encourage further high replication inside those cells.

Therefore, in order to create a more ’physiological’ model, we tried to establish another cellular study model in which the HT29 cells were used to mimic intestinal goblet cells. Notably, we had previously shown that the HT29 cell line is susceptible to SARS-CoV-2 infection but does not shed enough viral particles for the virus to be quantifiable in the cell culture media[24]. To achieve this, we co-cultured Caco-2 and HT29 cells (10:1 ratio) in order to form a single monolayer barrier as close as possible to the physiological conditions encountered in the mucosal epithelial layer containing enterocytes and goblet cells (**Fig. 6A)**. Under such conditions, the mucin produced by the HT29 cells (**Fig. S1**) should protect the Caco-2 cells, in a way similar to the protection of enterocytes in the intestinal mucosa.

**Figure 6.**
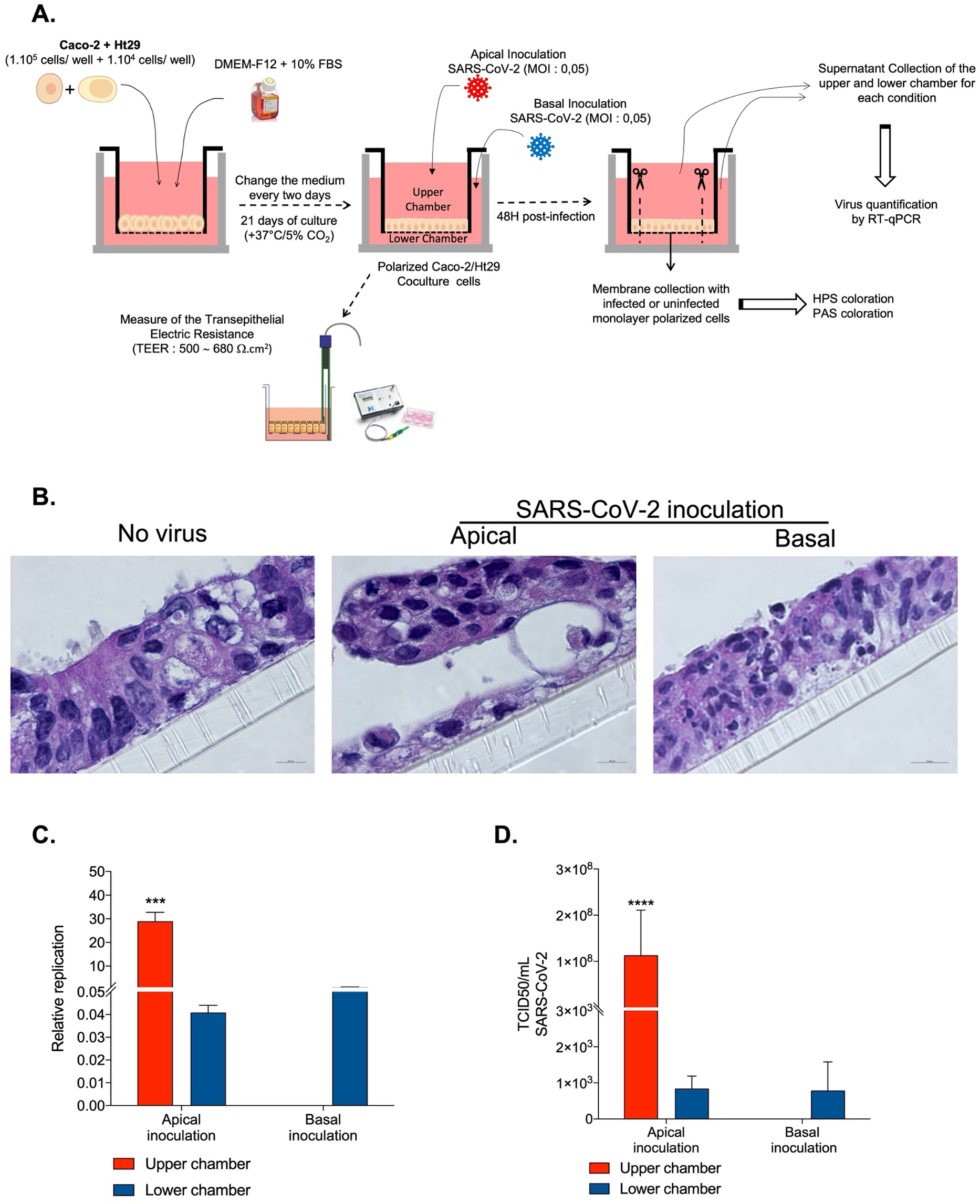
Impact of SARS-CoV-2 inoculation on the structure and permeability of the epithelial monolayer formed by the Caco-2/HT29 coculture. (**A**) Schematic flow of the analysis: After 21 days of culture in Transwell insert, the Caco-2/HT29 co-culture cells adopted an epithelial monolayer resembling an epithelial permeability barrier. The electronic trans-epithelial resistance (TEER) was measured during culture and was considered stable when it reached ∼500-700 Ω.cm^2^. Caco-2/HT29 cell monolayer were inoculated with the virus at apical (upper chamber) or basal side (lower chamber). At 48h post-infection, the polycarbonate membrane to which the monolayer cells adhere was collected for further microscopy analysis and the viral release was monitored from the cell’s supernatant from upper and lower chamber. (**B**) Microscopy analysis of the HPS coloration (Middle panel, scale bare: 10 μm, n=2) of the coculture inoculated with virus at apical side or basolateral side. (**C**) Quantification of viral release in the upper and lower chambers of the Caco-2/HT29 cell monolayer during apical inoculation and basolateral inoculation by qRT-PCR (n=2). The result was illustrated as ± SEM of the Relative quantification (RQ)= 2^-ΔCT^, where ΔCT represents the CT value obtained subtracted from the CT value at time T=0 (CT - CT_0_). T=0 corresponds to the viral CT at the precise moment of SARS-CoV-2 virus inoculation. (**D**) Quantification of SARS-CoV-2 infectious particles released in the supernatant of Caco-2/HT29 cells monolayer. TDCI50 (n=2) was performed by inoculating supernatants from the upper or lower chambers from SARS-CoV-2 infected Caco-2 cell monolayers into Vero E6 cell culture after 48h post-infection. The cytopathic effect (CPE) was evaluated on Vero E6 cells at 7days after exposure to the supernatants. The TCID50/mL is calculated according to the Spearman and Kärber algorithm.

As shown in **Fig. 6B** (left panel), in the absence of viral input, a dense and intact monolayer of polarised Caco-2/HT29 cells was observed by confocal microscopy analysis after HPS staining. At 48 hours post-inoculation of SARS-CoV-2 (MOI: 0.05) by the apical pole, evidence of less significant monolayer damage was observed (**Fig. 6B**, middle panel) compared to the previous model, since the cells were still adhering to the polycarbonate member. With basolateral virus inoculation, no major damage could be observed (**Fig. 6B**, right panel). As shown in **Fig. 6C** and **6D**, these histological changes specific to the apical inoculation are corroborated by the detection of virus at 48 hours post-inoculation in the lower chamber and a virus detection (viral input + progeny virus production) in the upper chamber corresponding to paracellular trafficking. In contrast, no virus could be detected in the upper chamber when the virus was inoculated at basolateral side. Although this model has some limitations, it provides direct evidence that the virus can create monolayer damage when inoculated at the apical pole, suggesting that in SARS-CoV-2 infected patients, the virus present in the lumen of the intestine might damage the intestinal epithelial barrier and reach the vasculature to spread to different organs.

## Discussion

Although the lungs are considered the primary site of SARS-CoV-2 replication, several studies have reported the presence of virus in the gastrointestinal tract [2]. One study found SARS-CoV-2 RNA on the hands of primary cases and contact cases, as well as on frequently-touched household surfaces associated with transmission, indicating these as potential vectors for spread within households[27]. We demonstrated that in K18-hACE2 transgenic mice infected with SARS-CoV-2 via intranasal administration, extensive necrosis and inflammation of the lamina propria of intestinal villi associated with cell damage and increased intestinal permeability are observed[24]. Very recently, iPSC-derived IEC monolayers have been shown to be suitable for SARS-CoV-2 research under physiologically relevant conditions[35].

In a Caco-2 cell intestinal cell line model, SARS-CoV-2 infection was found to depend on ACE2 and TMPRSS2[36] and is likely to occur through clathrin-mediated endocytosis[37,38]. Virions endocytosis into cells has been detected within minutes, and can be inhibited by neutralising antibodies to ACE2[39]. Although SARS-CoV-2 infects and replicates in Caco-2 cells, no clear cytopathic effect was detected in those cells[29,40]. In the polarised Caco-2 cell monolayer, the hepatitis A virus was reported to be mainly secreted in extracellular vesicles from the apical membrane (<1% from the basolateral membrane)[41]. Progeny virus release from the apical membrane of Caco-2 cells had also been reported with MERS-CoV[42]. However, more than two decades ago it was reported that polarised Caco-2 cells transfected with the murine Ceacam1 gene (a cellular receptor for the Mouse Hepatitis coronavirus, MHV), could be successfully infected with the MHV coronavirus, with progeny virions being released at the basolateral surface[43]. Interestingly, in a study on MERS-CoV infection of polarised intestinal cell monolayers, it was found that Caco-2 cells grown on Transwell inserts were more susceptible to viral infection upon inoculation of the virus at the apical than they were at the basolateral side, while inoculation of the virus into the stomach of a transgenic mouse expressing human DPP4 led to intestinal infection spreading to the respiratory tract and inducing animal death[42].

In an attempt to further explore this issue with SARS-CoV-2, we considered the development of a cellular model of polarised intestinal epithelial cells susceptible to SARS-CoV-2 as an appropriate tool to test the apical and/or basal route of infection of such cells. Thus, we developed a model of culture of Caco-2 cells on Transwell polycarbonate membranes, since we previously accumulated a huge amount of information regarding the infection of this cell line by SARS-CoV-2[24,29]. Recently we also reported that different SARS-CoV-2 isolates, namely B, B.1.416, B.1.367, B.1.1.7 (Alpha), B.1.351 (Beta), B.1.617.2 (Delta), P.1 (Gamma), A.27 and B.1.160, replicate similarly in Caco-2 cells[32] as well BA1.1 (Omicron) isolates[44]. However, none of these strains were isolated from faeces, they were all cultured in vitro from nasopharynx swabs. It should be emphasised that a higher genomic diversity of SARS-CoV-2 in faeces compared to the nasopharynx, highlighted a selection of viral strains that replicate in the gastrointestinal tract[45].

Here, we demonstrate that apical inoculation was much more effective in leading to virus (virus input + progeny virions) detection into the media in the upper and lower chambers than basolateral inoculation. Although we have not seen evidence of SARS-CoV-2 budding at the basolateral membrane, despite analysis of approximately one hundred sections of Caco-2 monolayers by electron microscopy, it remains possible that SARS-CoV-2 can undergo transcytosis across the epithelial barriers. However, it is much more likely that the virus we observed at the extracellular membrane on the apical side of polarised Caco-2 cells reach the bottom compartment of the Transwell after inducing damage in the Caco-2 cell barrier monolayer. Very similar results had been previously reported with MERS-CoV[42]. Histological examination revealed that apical SARS-CoV-2 inoculation (upper Transwell inoculation) triggers epithelial layer degeneration and live viruses emerged in the lower Transwell chamber, indicating that intercellular proteins junctions are damaged and no longer act as a barrier. Furthermore, there are numerous pieces of evidence that SARS-CoV-2 infection disrupts tight junctions by metallopeptidases in the intestinal epithelium and Caco-2 monolayer[2,24,46,47] crossing the intestinal epithelial barriers by paracellular trafficking, as previously reported with other viruses[48–50]. It is also worth noting that the cellular receptor for SARS-CoV, angiotensin-converting enzyme 2 (ACE2), is located on the apical plasma membrane of polarized epithelial cells and mediates infection from the apical side of these cells[51].

In contrast to Caco-2 cells, HT29 cells are poorly susceptible to SARS-CoV-2 replication and do not sustain viral production[24,29] . This lack of viral production was unlikely to be due to the low expression of mRNA ACE2 [24], as well a low expression of the receptor on the surface of these cells[52]. However, this low expression could be linked to the expression of membrane-associated, glycan-dependent mucins[53,54], which may exhibit potent anti-SARS-CoV-2 activity. Interestingly, when polarised Caco-2/HT29 cell monolayers were tested for virus detection in the upper and lower chambers of the Transwell insert, the presence of virus was observed in both chambers after apical inoculation, but in the lower chamber only after basolateral inoculation. Furthermore, in comparison with the previous model, i.e. polarised Caco-2 alone, the concentration of SARS-CoV-2 found in the lower chamber during apical inoculation was considerably higher than in the co-culture model. We also found that the integrity of the monolayer was less damaged after apical virus inoculation, since the cells were still adhering to the polycarbonate membrane. This suggests that the cell barrier composed by polarised Caco-2/HT29 cell monolayers was possibly more watertight than monolayers composed of Caco-2 cells only.

Although our model may provide interesting information, suggesting that in patients infected with SARS-CoV-2 the virus present in the lumen of the intestine might damage the intestinal epithelial barrier and reach the vasculature to spread into different organs, it remains simplified compared to the microenvironment that the virus will encounter into the gastrointestinal tract of infected patients. In patients infected with SARS-CoV-2, IgA molecules synthesised by immune cells in the lamina propria are likely to predominate the secretions of intestinal mucous membranes, promoting virus transcytosis towards the apical membranes after interaction with the polymeric immunoglobulin receptor[55]. IgG levels can also significantly increase in response to SARS-CoV-2 infection and control the infection process. SARS-CoV-2 infection induces a change in the gut immunological barrier[56,57].

Since several studies suggest that the gastrointestinal tract may be an entry portal and also a reservoir for long COVID, our results provide information on the dynamic of intestinal cell monolayer infection by SARS-CoV-2 with evidence of predominant apical versus basal infection of Caco-2 and Caco-2/HT29 cell monolayers by SARS-CoV-2. Actually, our results are supported by other studies carried out by our research peers on polarised airway epithelial cells in which they found very similar results, namely the efficacy of both SARS-CoV[58] and SARS-CoV-2[59,60] infection via the apical pole leading to damage of the epithelial barrier.

Although we cannot exclude trancytosis events, our results suggest a preferential disruption of tight junctions, damage of monolayer barrier integrity, and paracellular trafficking of the virus.

## Acknowledgments

We thank Prof. J-C Lagier, Prof. D Raoult and Prof. B La Scola for stimulating discussions and unwavering support. We thank Dr. J-P Baudoin for his expert advice on electron microscopy. We thank Prof. H Lepidi for allowing us access to the equipment of the anatomopathology platform at the La Timone hospital (APHM, Marseille), and Prof. B La Scola for allowing us access to the BSL3 facility of the IHU-Méditerranée Infection.

## Authors Contributions

IOO and CD contributed to the design of the study and conceived the manuscript. CG and IO performed the viral infections and the in vitro experiments. JA and JA performed the electron microscopy analysis. J-LM, BD and CD supervised the work. CD wrote the first draft of the manuscript. All authors participated in correction of the manuscript. All authors contributed to the article and approved the submitted version.

## Figure Legends

**Figure S1.**
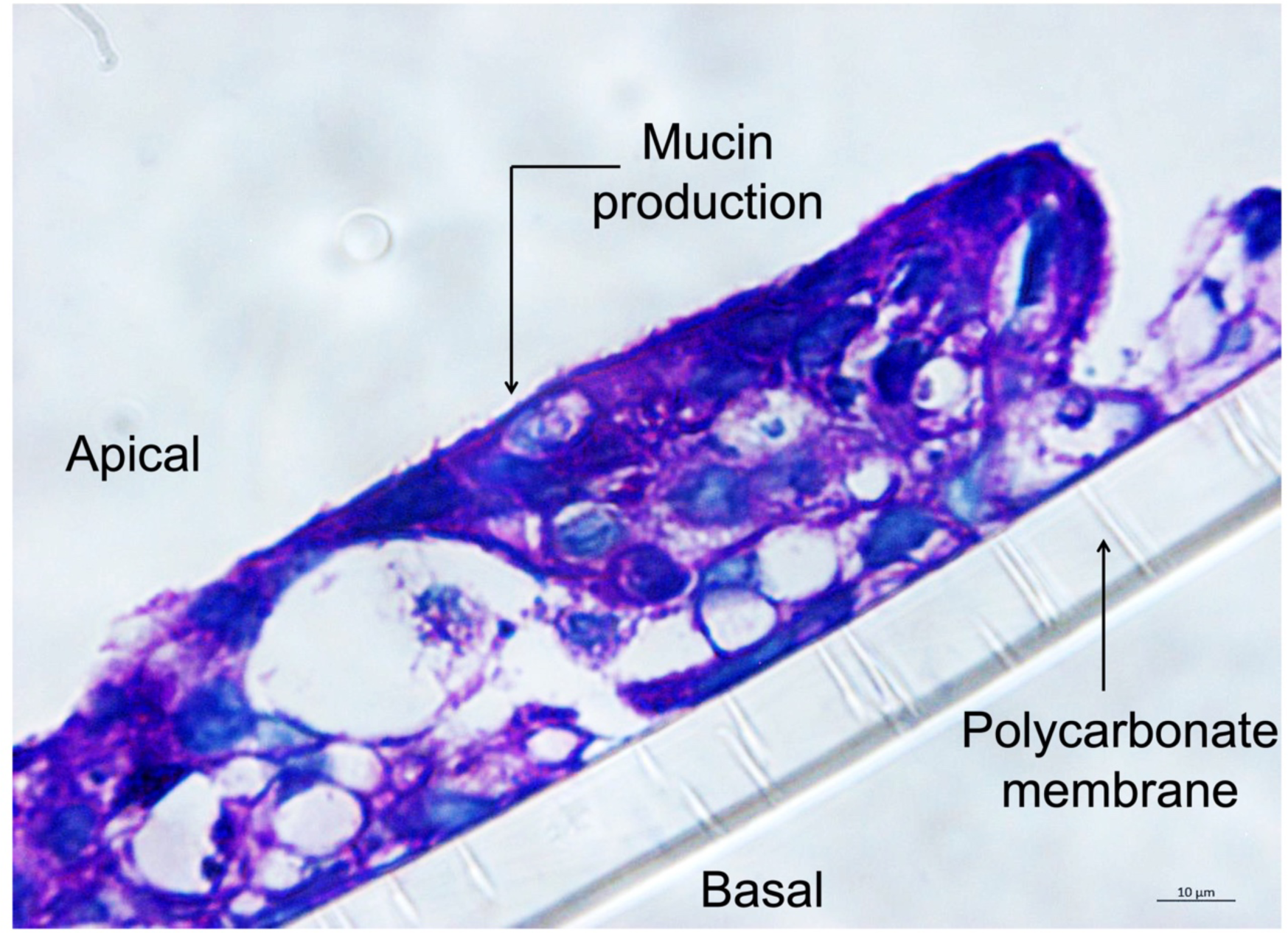
Microscopy analysis of the PAS (Periodic Acid-Schiff) coloration for the mucin detection in the Caco-2/HT29 coculture. Scale bare: 10 μm, n=2.

